# Connecting the dots between mechanosensitive channel abundance, osmotic shock, and survival at single-cell resolution

**DOI:** 10.1101/339259

**Authors:** Griffin Chure, Heun Jin Lee, Rob Phillips

## Abstract

Rapid changes in extracellular osmolarity are one of many insults microbial cells face on a daily basis. To protect against such shocks, *Escherichia coli* and other microbes express several types of transmembrane channels which open and close in response to changes in membrane tension. In *E. coli*, one of the most abundant channels is the mechanosensitive channel of large conductance (MscL). While this channel has been heavily characterized through structural methods, electrophysiology, and theoretical modeling, our understanding of its physiological role in preventing cell death by alleviating high membrane tension remains tenuous. In this work, we examine the contribution of MscL alone to cell survival after osmotic shock at single cell resolution using quantitative fluorescence microscopy. We conduct these experiments in an *E. coli* strain which is lacking all mechanosensitive channel genes save for MscL whose expression is tuned across three orders of magnitude through modifications of the Shine-Dalgarno sequence. While theoretical models suggest that only a few MscL channels would be needed to alleviate even large changes in osmotic pressure, we find that between 500 and 700 channels per cell are needed to convey upwards of 80% survival. This number agrees with the average MscL copy number measured in wild-type *E. coli* cells through proteomic studies and quantitative Western blotting. Furthermore, we observe zero survival events in cells with less than 100 channels per cell. This work opens new questions concerning the contribution of other mechanosensitive channels to survival as well as regulation of their activity.

## Importance

Mechanosensitive (MS) channels are transmembrane protein complexes which open and close in response to changes in membrane tension as a result of osmotic shock. Despite extensive biophysical characterization, the contribution of these channels to cell survival remains largely unknown. In this work, we use quantitative video microscopy to measure the abundance of a single species of MS channel in single cells followed by their survival after a large osmotic shock. We observe total death of the population with less than 100 channels per cell and determine that approximately 500 – 700 channels are needed for 80% survival. The number of channels we find to confer nearly full survival is consistent with the counts of the number of channels in wild type cells in several earlier studies. These results prompt further studies to dissect the contribution of other channel species to survival.

## Introduction

Changes in the extracellular osmolarity can be a fatal event for the bacterial cell. Upon a hypo-osmotic shock, water rushes into the cell across the membrane, leaving the cell with no choice but to equalize the pressure. This equalization occurs either through damage to the cell membrane (resulting in death) or through the regulated flux of water molecules through transmembrane protein channels (Fig 1A). Such proteinaceous pressure release valves have been found across all domains of life, with the first bacterial channel being described in 1987 (1). Over the past thirty years, several more channels have been discovered, described, and (in many cases) biophysically characterized. *E. coli*, for example, has seven of these channels (one MscL and six MscS homologs) which have varied conductance, gating mechanisms, and expression levels. While they have been the subject of much experimental and theoretical dissection, much remains a mystery with regard to the roles their abundance and interaction with other cellular processes play in the greater context of physiology (2–8).

**FIG 1.**
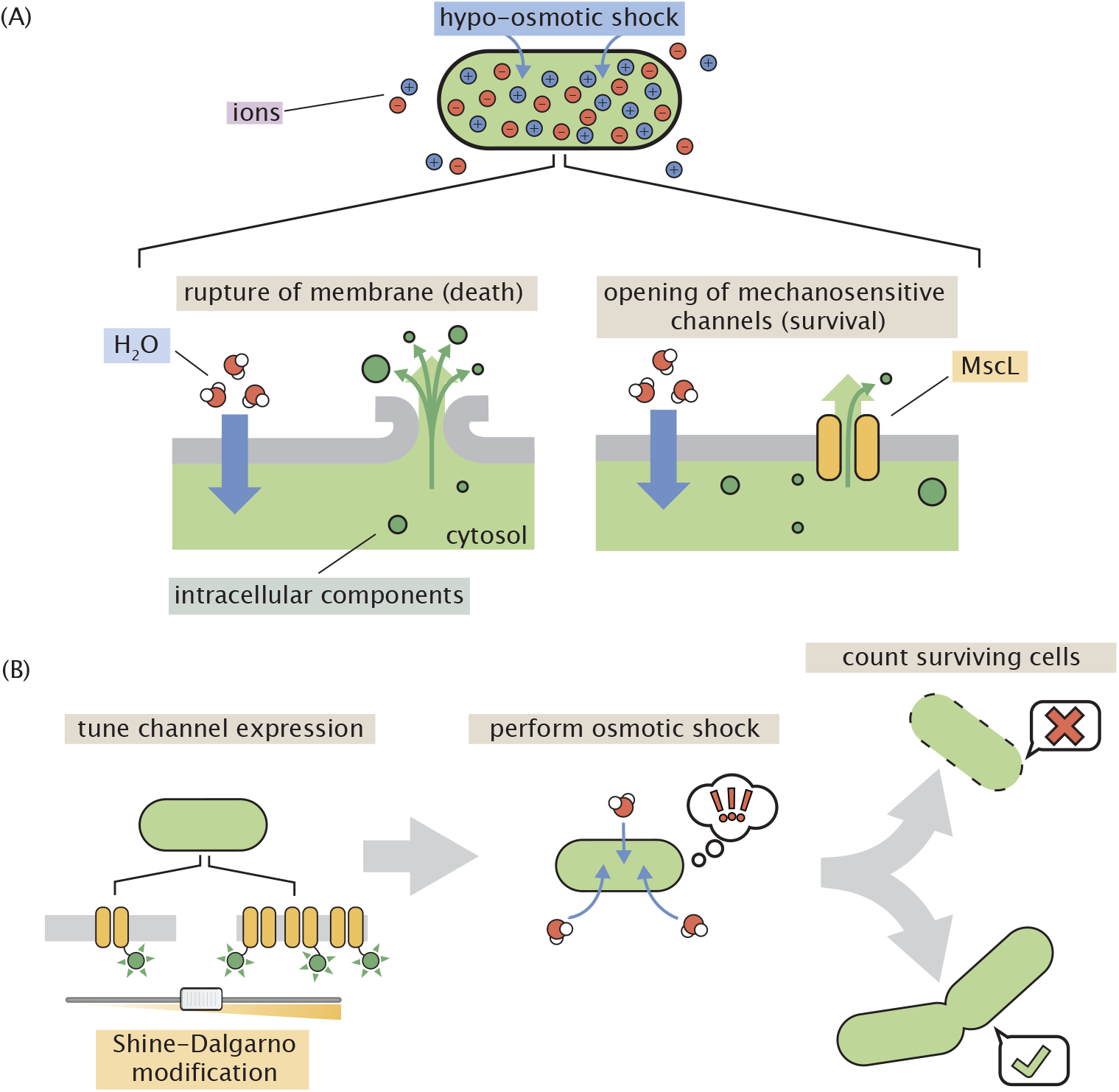
Role of mechanosensitive channels during hypo-osmotic shock. (A) A hypo-osmotic shock results in a large difference in the osmotic strength between the intracellular and extracellular spaces. As a result, water rushes into the cell to equalize this gradient increasing the turgor pressure and tension in the cell membrane. If no mechanosensitive channels are present and membrane tension is high (left panel), the membrane ruptures releasing intracellular content into the environment resulting in cell death. If mechanosensitive channels are present (right panel) and membrane tension is beyond the gating tension, the mechanosensitive channel MscL opens, releasing water and small intracellular molecules into the environment thus relieving pressure and membrane tension. (B) The experimental approach undertaken in this work. The number of mechanosensitive channels tagged with a fluorescent reporter is tuned through modification of the Shine-Dalgarno sequence of the *mscL* gene. The cells are then subjected to a hypo-osmotic shock and the number of surviving cells are counted, allowing the calculation of a survival probability.

Of the seven channels in *E. coli*, the mechanosensitive channel of large conductance (MscL) is one of the most abundant and the best characterized. This channel has a large conductance (3 nS) and mediates the flux of water molecules across the membrane via a ~3 nm wide pore in the open state (9, 10). Molecular dynamics simulations indicate that a single open MscL channel permits the flux of 4 × 10^9^ water molecules per second, which is an order of magnitude larger than a single aquaporin channel (BNID 100479) (11, 12). This suggests that having only a few channels per cell could be sufficient to relieve even large changes in membrane tension. Electrophysiological experiments have suggested a small number of channels per cell (13, 14), however, more recent approaches using quantitative western blotting, fluorescence microscopy, and proteomics have measured several hundred MscL per cell (3, 15, 16). To further complicate matters, the expression profile of MscL appears to depend on growth phase, available carbon source, and other environmental challenges (3, 16, 17). While there are likely more than just a few channels per cell, why cells seem to need so many and the biological rationale behind their condition-dependent expression both remain a mystery.

While their biochemical and biophysical characteristics have received much attention, their connection to cell survival is understudied. Drawing such a direct connection between channel copy number and survival requires quantitative *in vivo* experiments. To our knowledge, the work presented in van den Berg et al. 2016 (8) is the first attempt to simultaneously measure channel abundance and survivability for a single species of mechanosensitive channel. While the measurement of channel copy number was performed at the level of single cells using super-resolution microscopy, survivability after a hypo-osmotic shock was assessed in bulk plating assays which rely on serial dilutions of a shocked culture followed by counting the number of resulting colonies after incubation. Such bulk assays have long been the standard for querying cell viability after an osmotic challenge. While they have been highly informative, they reflect only the mean survival rate of the population, obfuscating the variability in survival of the population. The stochastic nature of gene expression results in a noisy distribution of MscL channels rather than a single value, meaning those found in the long tails of the distribution have quite different survival rates than the mean but are lost in the final calculation of survival probability.

In this work, we present an experimental system to quantitatively probe the interplay between MscL copy number and survival at single-cell resolution, as is seen in Fig. 1B. We generated an *E. coli* strain in which all seven mechanosensitive channels had been deleted from the chromosome followed by a chromosomal integration of a single gene encoding an MscL-super-folder GFP (sfGFP) fusion protein. To explore copy number regimes beyond those of the wild-type expression level, we modified the Shine-Dalgarno sequence of this integrated construct allowing us to cover nearly three decades of MscL copy number. To probe survivability, we exposed cells to a large hypo-osmotic shock at controlled rates in a flow cell under a microscope, allowing the observation of the single-cell channel copy number and the resulting survivability of single cells. With this large set of single cell measurements, we approach the calculation of survival probability in a manner that is free of binning bias which allows the reasonable extrapolation of survival probability to copy numbers outside of the observed range. In addition, we show that several hundred channels are needed to convey high rates of survival and observe a minimum number of channels needed to permit any degree of survival.

## Results

### Quantifying the single-cell MscL copy number

The principal goal of this work is to examine the contribution of a single mechanosensitive channel species to cell survival under a hypo-osmotic shock. While this procedure could be performed for any species of channel, we chose MscL as it is the most well characterized and one of the most abundant species in *E. coli*. To probe the contribution of MscL alone, we generated an *E. coli* strain in which all seven known mechanosensitive channel genes were deleted from the chromosome followed by the integration of an *mscL* gene encoding an MscL super-folder GFP (sfGFP) fusion. Chromosomal integration imposes strict control on the gene copy number compared to plasmid borne expression systems, which is important to minimize variation in channel expression across the population and provide conditions more representative of native cell physiology. Fluorescent protein fusions have frequently been used to study MscL and have been shown through electrophysiology to function identically to the native MscL protein, allowing us to confidently draw conclusions about the role this channel plays in wild-type cells from our measurements. (3, 18).

To modulate the number of MscL channels per cell, we developed a series of mutants which were designed to decrease the expression relative to wild-type. These changes involved direct alterations of the Shine-Dalgarno sequence as well as the inclusion of AT hairpins of varying length directly upstream of the start codon which influences the translation rate and hence the number of MscL proteins produced (Fig. 2A). The six Shine-Dalgarno sequences used in this work were chosen using the RBS binding site strength calculator from the Salis Laboratory at the Pennsylvania State University (19, 20). While the designed Shine-Dalgarno sequence mutations decreased the expression relative to wild-type as intended, the distribution of expression is remarkably wide spanning an order of magnitude.

**FIG 2.**
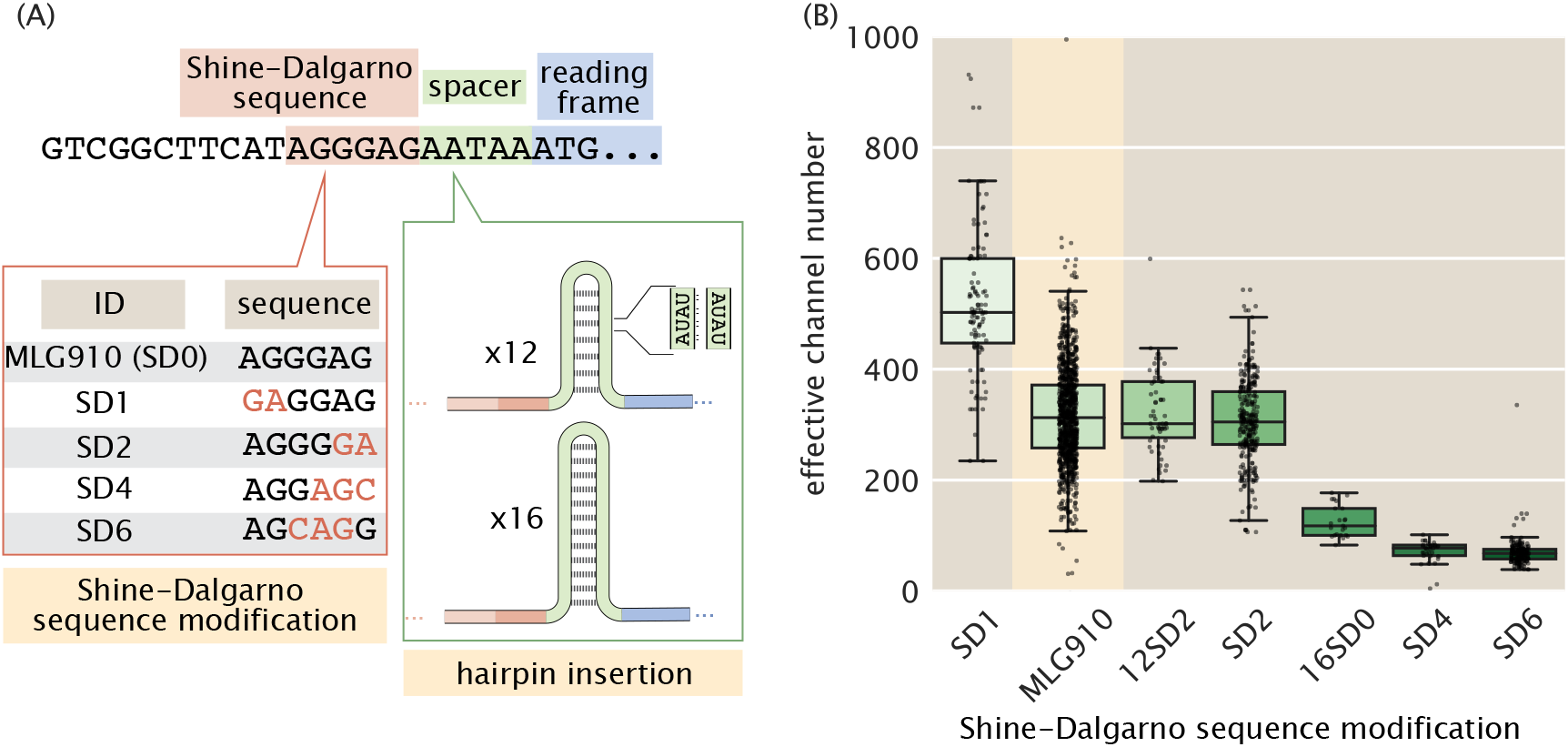
Control of MscL expression and calculation of channel copy number. (A) Schematic view of the expression modifications performed in this work. The beginning portion of the native *mscL* sequence is shown with the Shine-Dalgarno sequence, spacer region, and start codon shaded in red, green, and blue, respectively. The Shine-Dalgarno sequence was modified through the Salis lab Ribosomal Binding Strength calculator (19, 20). The wild-type sequence (MLG910) is shown in black with mutations for the other four Shine-Dalgarno mutants highlighted in red. Expression was further modified by the insertion of repetitive AT bases into the spacer region, generating hairpins of varying length which acted as a thermodynamic barrier for translation initiation. (B) Variability in effective channel copy number is computed using the standard candle. The boxes represent the interquartile region of the distribution, the center line displays the median, and the whiskers represent 1.5 times the maximum and minimum of the interquartile region. Individual measurements are denoted as black points. The strain used for calibration of channel copy number (MLG910) is highlighted in yellow.

To measure the number of MscL channels per cell, we determined a fluorescence calibration factor to translate arbitrary fluorescence units per cell to protein copy number. While there have been numerous techniques developed over the past decade to directly measure this calibration factor, such as quantifying single-molecule photobleaching constants or measuring the binomial partitioning of fluorescent proteins upon cell division (3, 21), we used *a priori* knowledge of the mean MscL-sfGFP expression level of a particular *E. coli* strain to estimate the average fluorescence of a single channel. In Bialecka-Fornal et al. 2012 (3), the authors used single-molecule photobleaching and quantitative Western blotting to probe the expression of MscL-sfGFP under a wide range of growth conditions. To compute a calibration factor, we used the strain MLG910 (*E. coli* K12 MG1655 *ϕ*(mscL-sfGFP)) as a “standard candle”, highlighted in yellow in Fig. 2B. This standard candle strain was grown and imaged in identical conditions in which the MscL count was determined. The calibration factor was computed by dividing the mean total cell fluorescence by the known MscL copy number, resulting in a measure of arbitrary fluorescence units per MscL channel. Details regarding this calculation and appropriate propagation of error can be found in the Materials & Methods as well as the supplemental information (*Standard Candle Calibration*).

While it is seemingly trivial to use this calibration to determine the total number of channels per cell for wild-type or highly expressing strains, the calculation for the lowest expressing strains is complicated by distorted cell morphology. We observed that as the channel copy number decreases, cellular morphology becomes increasingly aberrant with filamentous, bulging, and branched cells becoming more abundant (Fig. S3A). This morphological defect has been observed when altering the abundance of several species of mechanosensitive channels, suggesting that they play an important role in general architectural stability (3, 4). As these aberrant morphologies can vary widely in size and shape, calculating the number of channels per cell becomes a more nuanced endeavor. For example, taking the total MscL copy number for these cells could skew the final calculation of survival probability as a large but severely distorted cell would be interpreted as having more channels than a smaller, wild-type shaped cell (Fig. S3B). To correct for this pathology, we computed the average expression level per unit area for each cell and multiplied this by the average cellular area of our standard candle strain which is morphologically indistinguishable from wild-type *E. coli*, allowing for the calculation of an effective channel copy number. The effect of this correction can be seen in Fig. S3C and D, which illustrate that there is no other correlation between cell area and channel expression.

Our calculation of the effective channel copy number for our suite of Shine-Dalgarno mutants is shown in Fig. 2B. The expression of these strains cover nearly three orders of magnitude with the extremes ranging from approximately four channels per cell to nearly one thousand. While the means of each strain are somewhat distinct, the distributions show a large degree of overlap, making one strain nearly indistinguishable from another. This variance is a quantity that is lost in the context of bulk scale experiments but can be accounted for via single-cell methods.

### Performing a single-cell hypo-osmotic challenge assay

To measure the channel copy number of a single cell and query its survival after a hypo-osmotic shock, we used a custom-made flow cell in which osmotic shock and growth can be monitored in real time using video microscopy (Fig. 3A). The design and characterization of this device has been described in depth previously and is briefly described in the Materials & Methods (4). Using this device, cells were exposed to a large hypo-osmotic shock by switching between LB Miller medium containing 500mM NaCl and LB media containing no NaCl. All six Shine-Dalgarno modifications shown in Fig. 2B (excluding MLG910) were subjected to a hypo-osmotic shock at controlled rates while under observation. After the application of the osmotic shock, the cells were imaged every sixty seconds for four to six hours. Survivors were defined as cells which underwent at least two divisions post-shock. The brief experimental protocol can be seen in Fig. 3B.

**FIG 3.**
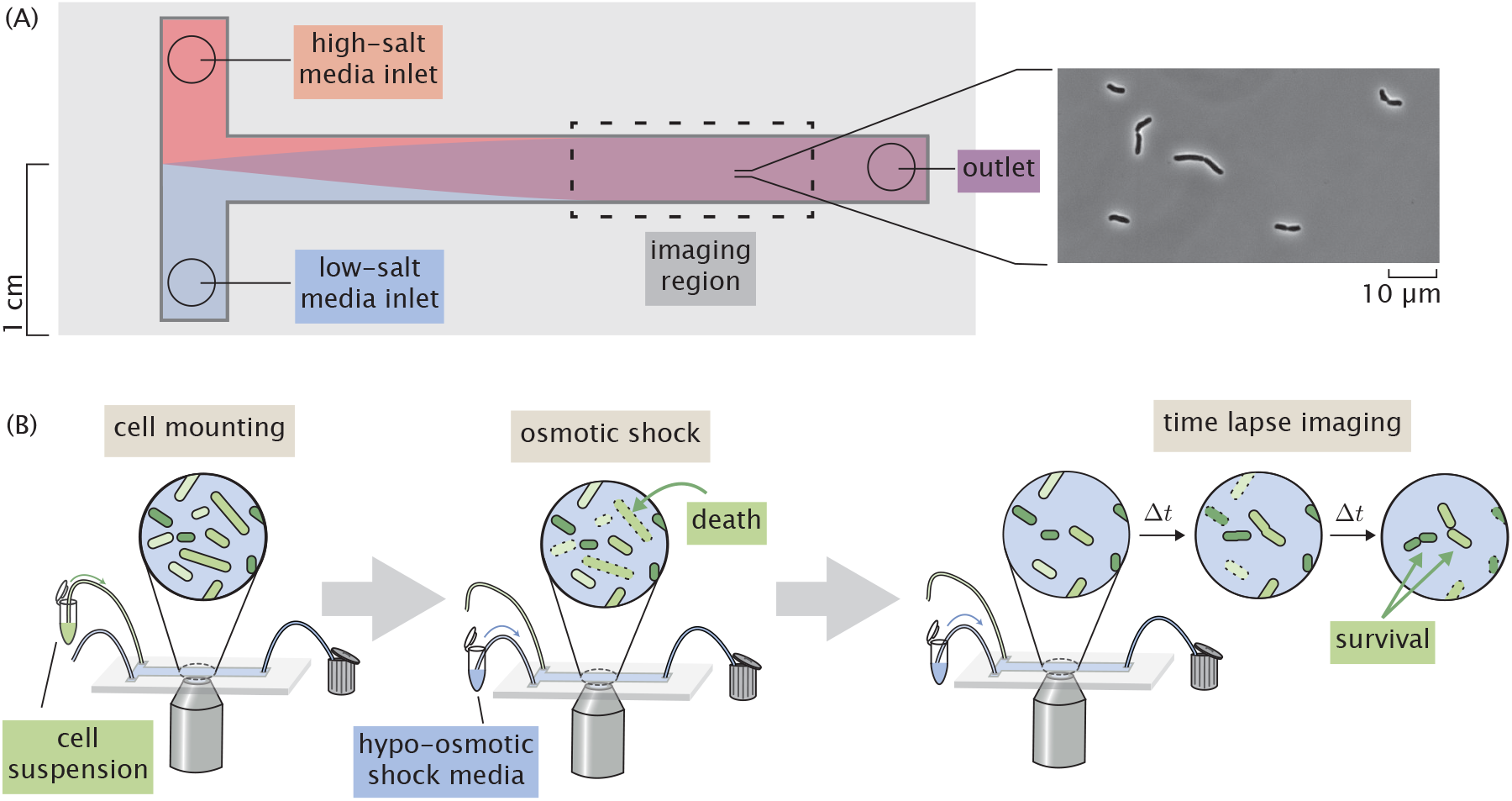
Experimental approach to measuring survival probability. (A) Layout of a home-made flow cell for subjecting cells to osmotic shock. Cells are attached to a polyethylamine functionalized surface of a glass coverslip within the flow chamber by loading a dilute cell suspension through one of the inlets. (B) The typical experimental procedure. Cells are loaded into a flow chamber as shown in (A) and mounted to the glass coverslip surface. Cells are subjected to a hypo-osmotic shock by flowing hypotonic medium into the flow cell. After shock, the cells are monitored for several hours and surviving cells are identified.

Due to the extensive overlap in expression between the different Shine-Dalgarno mutants (see Fig. 2B), computing the survival probability by treating each mutant as an individual bin obfuscates the relationship between channel abundance and survival. To more thoroughly examine this relationship, all measurements were pooled together with each cell being treated as an individual experiment. The hypo-osmotic shock applied in these experiments was varied across a range of 0.02 Hz (complete exchange in 50 s) to 2.2 Hz (complete exchange in 0.45 s). Rather than pooling this wide range of shock rates into a single data set, we chose to separate the data into “slow shock” (< 1.0 Hz) and “fast shock” (≥ 1.0 Hz) classes. Other groupings of shock rate were explored and are discussed in the supplemental information (*Shock Classification*). The cumulative distributions of channel copy number separated by survival are shown in Fig. 4. In these experiments, survival was never observed for a cell containing less than approximately 100 channels per cell, indicated by the red stripe in Fig. 4. This suggests that there is a minimum number of channels needed for survival on the order of 100 per cell. We also observe a slight shift in the surviving fraction of the cells towards higher effective copy number, which matches our intuition that including more mechanosensitive channels increases the survival probability.

**FIG 4.**
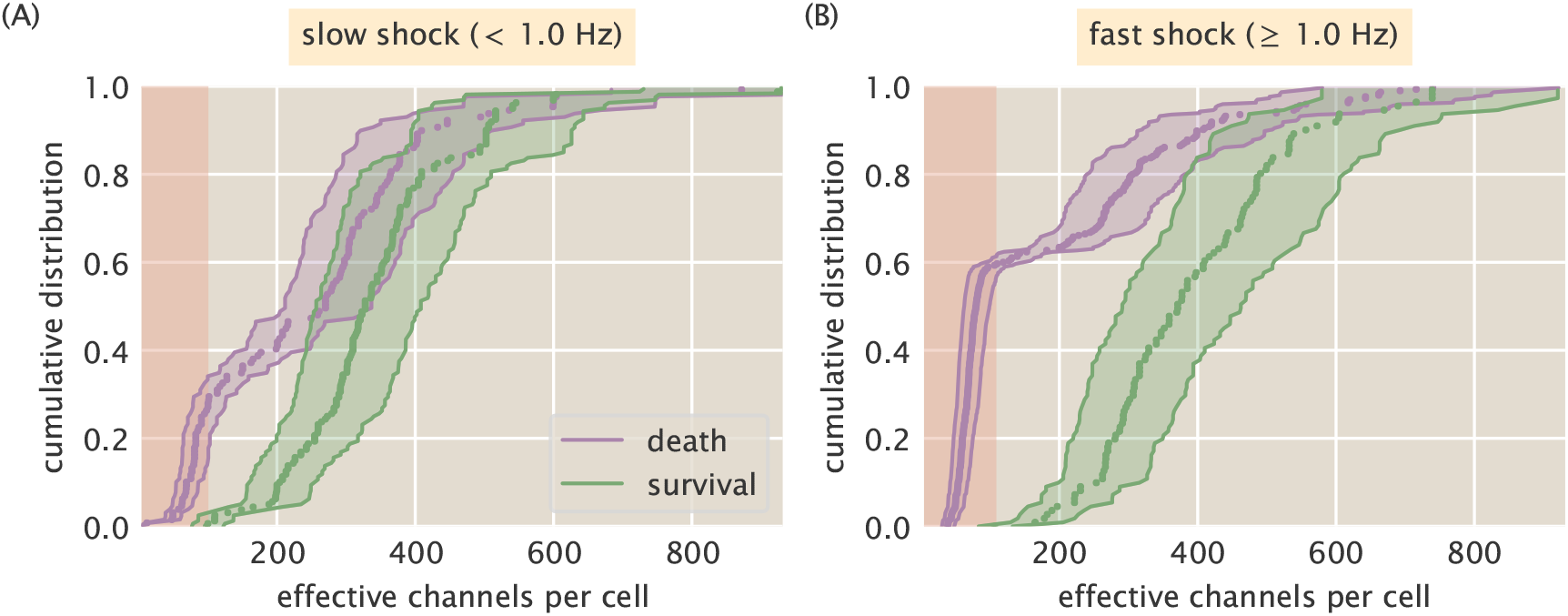
Distributions of survival and death as a function of effective channel number. (A) Empirical cumulative distributions of channel copy number separated by survival (green) or death (purple) after a slow (< 1.0 Hz) osmotic shock. (B) The empirical cumulative distribution for a fast (≥ 1.0 Hz) osmotic shock. Shaded green and purple regions represent the 95% credible region of the effective channel number calculation for each cell. Shaded red stripe signifies the range of channels in which no survival was observed.

### Prediction of survival probability as a function of channel copy number

There are several ways by which the survival probability can be calculated. The most obvious approach would be to group each individual Shine-Dalgarno mutant as a single bin and compute the average MscL copy number and the survival probability. Binning by strain is the most frequently used approach for such measurements and has provided valuable insight into the qualitative relationship of survival on other physiological factors (4, 8). However the copy number distribution for each Shine-Dalgarno mutant (Fig. 2B) is remarkably wide and overlaps with the other strains. We argue that this coarse-grained binning negates the benefits of performing single-cell measurements as two strains with different means but overlapping quartiles would be treated as distinctly different distributions.

Another approach would be to pool all data together, irrespective of the Shine-Dalgarno mutation, and bin by a defined range of channels. Depending on the width of the bin, this could allow for finer resolution of the quantitative trend, but the choice of the bin width is arbitrary with the *a priori* knowledge that is available. Drawing a narrow bin width can easily restrict the number of observed events to small numbers where the statistical precision of the survival probability is lost. On the other hand, drawing wide bins increases the precision of the estimate, but becomes further removed from a true single-cell measurement and represents a population mean, even though it may be a smaller population than binning by the Shine-Dalgarno sequence alone. In both of these approaches, it is difficult to extrapolate the quantitative trend outside of the experimentally observed region of channel copy number. Here, we present a method to estimate the probability of survival for any channel copy number, even those that lie outside of the experimentally queried range.

To quantify the survival probability while maintaining single-cell resolution, we chose to use a logistic regression model which does not require grouping data into arbitrary bins and treats each cell measurement as an independent experiment. Logistic regression is an inferential method to model the probability of a boolean or categorical event (such as survival or death) given one or several predictor variables and is commonly used in medical statistics to compute survival rates and dose response curves (22, 23). The primary assumption of logistic regression is that the log-odds probability of survival *p_s_* is linearly dependent on the predictor variable, in our case the log channels per cell *N_c_* with a dimensionless intercept *β*_0_ and slope *β*_1_,

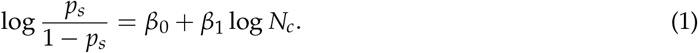

Under this assumption of linearity, *β*_0_ is the log-odds probability of survival with no MscL channels. The slope *β*_1_ represents the change in the log-odds probability of survival conveyed by a single channel. As the calculated number of channels in this work spans nearly three orders of magnitude, it is better to perform this regression on log *N_c_* as regressing on *N_c_* directly would give undue weight for lower channel copy numbers due to the sparse sampling of high-copy number cells. The functional form shown in Eq. 1 can be derived directly from Bayes’ theorem and is shown in the supplemental information (*Logistic Regression*). If one knows the values of *β*_0_ and *β*_1_, the survival probability can be expressed as

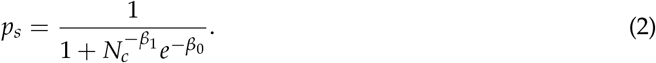

In this analysis, we used Bayesian inferential methods to determine the most likely values of the coefficients and is described in detail in the supplemental information (*Logistic Regression*).

The results of the logistic regression are shown in Fig. 5. We see a slight rightward shift the survival probability curve under fast shock relative to the slow shock case, reaffirming the conclusion that survival is also dependent on the rate of osmotic shock (4). This rate dependence has been observed for cells expressing MscL alongside other species of mechanosensitive channels, but not for MscL alone. This suggests that MscL responds differently to different rates of shock, highlighting the need for further study of rate dependence and the coordination between different species of mechanosensitive channels. Fig. 5 also shows that several hundred channels are required to provide appreciable protection from osmotic shock. For a survival probability of 80%, a cell must have approximately 500 to 700 channels per cell for a fast and slow shock, respectively. The results from the logistic regression are showed as continuous colored curves. The individual cell measurements separated by survival and death are shown at the top and bottom of each plot, respectively, and are included to provide a sense of sampling density.

**FIG 5.**
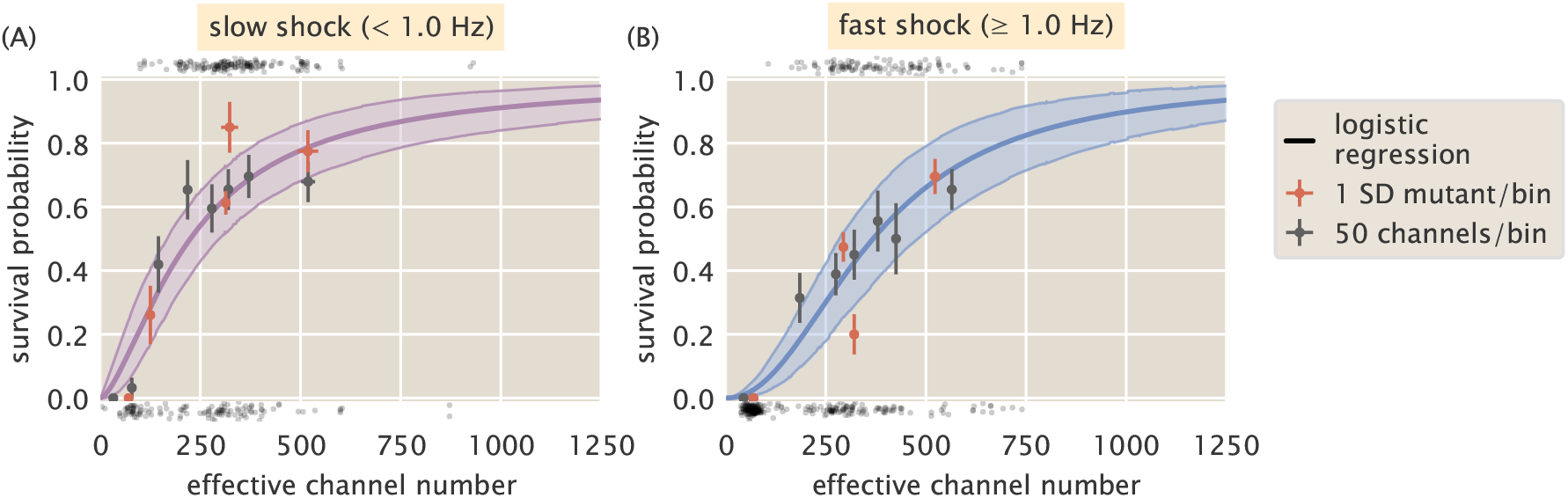
Probability of survival as a function of MscL copy number. (A) Estimated survival probability for survival under slow shock as a function of channel copy number. (B) The estimated survival probability of survival under a fast shock as a function of channel copy number. Solid curves correspond to the most probable survival probability from a one-dimensional logistic regression. Shaded regions represent the 95% credible regions. Points at the top and bottom of plots represent individual cell measurements which survived and perished, respectively. The red and black points correspond to the survival probability estimated via binning by Shine-Dalgarno sequence and binning by groups of 50 channels per cell, respectively. Horizontal error bars represent the standard error of the mean from at least 25 measurements. Vertical error bars represent the certainty of the probability estimate given *n* survival events from *N* total observations.

Over the explored range of MscL copy number, we observed a maximum of 80% survival for any binning method. The remaining 20% survival may be attained when the other species of mechanosensitive channels are expressed alongside MscL. However, it is possible that the flow cell method performed in this work lowers the maximal survival fraction as the cells are exposed to several, albeit minor, mechanical stresses such as loading into the flow cell and chemical adherence to the glass surface. To ensure that the results from logistic regression accurately describe the data, we can compare the survival probabilities to those using the binning methods described earlier (red and black points, Fig. 5). Nearly all binned data fall within error of the prediction (see Materials & Methods for definition of error bar on probability), suggesting that this approach accurately reflects the survival probability and gives license to extrapolate the estimation of survival probability to regions of outside of our experimentally explored copy number regime.

Thus far, we’ve dictated that for a given rate of osmotic shock (i.e. “fast” or “slow”), the survival probability is dependent only on the number of channels. In Fig. S7, we show the result of including other predictor variables, such as area and shock rate alone. In such cases, including other predictors resulted in pathological curves showing that channel copy number is the most informative out of the available predictor variables.

## Discussion

One of the most challenging endeavors in the biological sciences is linking the microscopic details of cellular components to the macro-scale physiology of the organism. This formidable task has been met repeatedly in the recent history of biology, especially in the era of DNA sequencing and single molecule biochemistry. For example, the scientific community has been able to connect sickle-cell anemia to a single amino acid substitution in Hemoglobin which promotes precipitation under a change in O_2_ partial pressure (24–26). Others have assembled a physical model that quantitatively describes chemosensation in bacteria (27) in which the arbiter of sensory adaptation is the repeated methylation of chemoreceptors (28–31). In the past ~50 years alone, numerous biological and physical models of the many facets of the central dogma have been assembled that give us a sense of the interplay between the genome and physiology. For example, the combination of biochemical experimentation and biophysical models have given us a picture of how gene dosage affects furrow positioning in *Drosophila* (32), how recombination of V(D)J gene segments generates an extraordinarily diverse antibody repertoire (33–35), and how telomere shortening through DNA replication is intrinsically tied to cell senescence (36, 37), to name just a few of many such examples.

By no means are we “finished” with any of these topics. Rather, it’s quite the opposite in the sense that having a handle on the biophysical knobs that tune the behavior opens the door to a litany of new scientific questions. In the case of mechanosenstaion and osmoregulation, we have only recently been able to determine some of the basic facts that allow us to approach this fascinating biological phenomenon biophysically. The dependence of survival on mechanosensitive channel abundance is a key quantity that is missing from our collection of critical facts. To our knowledge, this work represents the first attempt to quantitatively control the abundance of a single species of mechanosensitive channel and examine the physiological consequences in terms of survival probability at single-cell resolution. Our results reveal two notable quantities. First, out of the several hundred single-cell measurements, we never observed a cell which had less than approximately 100 channels per cell and survived an osmotic shock, irrespective of the shock rate. The second is that between 500 and 700 channels per cell are needed to provide ≥ 80% survival, depending on the shock rate.

Only recently has the relationship between the MscL copy number and the probability of survival been approached experimentally. In van den Berg et al. (2016), the authors examined the contribution of MscL to survival in a genetic background where all other known mechanosensitive channels had been deleted from the chromosome and plasmid-borne expression of an MscL-mEos3.2 fusion was tuned through an IPTG inducible promoter (8). In this work, they measured the single-cell channel abundance through super-resolution microscopy and queried survival through bulk assays. They report a nearly linear relationship between survival and copy number, with approximately 100 channels per cell conveying 100% survival. This number is significantly smaller than our observation of approximately 100 channels as the *minimum* number needed to convey any observable degree of survival.

The disagreement between the numbers reported in this work and in van den Berg et al. may partially arise from subtle differences in the experimental approach. The primary practical difference is the rate and magnitude of the osmotic shock. van den Berg et al. applied an approximately 600 mOsm downshock in bulk at an undetermined rate whereas we applied a 1 Osm downshock at controlled rates varying from 0.02 Hz to 2.2 Hz. In their work, survival was measured through plating assays which represent the population average rather than the distribution of survival probability. While this approach provides valuable information regarding the response of a population to an osmotic shock, the high survival rate may stem from a wide distribution of channel copies per cell in the population coupled with bulk-scale measurement of survival. As has been shown in this work, the expression of MscL from a chromosomal integration is noisy with a single strain exhibiting MscL copy numbers spanning an order of magnitude or more. In van den Berg et al., this variance is exacerbated by expression of MscL from an inducible plasmid as fluctuations in the gene copy number from plasmid replication and segregation influence the expression level. Connecting such a wide and complex distribution of copy numbers to single-cell physiology requires the consideration of moments beyond the mean which is a nontrivial task. Rather than trying to make such a connection, we queried survival at single-cell resolution at the expense of a lower experimental throughput.

Despite these experimental differences, the results of this work and van den Berg et al., are in agreement that MscL must be present at the level of 100 or more channels per cell in wild-type cells to convey appreciable survival. As both of these works were performed in a strain in which the only mechanosensitive channel was MscL, it remains unknown how the presence of the other channel species would alter the number of MscL needed for complete survival. In our experiments, we observed a maximum survival probability of approximately 80% even with close to 1000 MscL channels per cell. It is possible that the combined effort of the six other mechanosensitive channels would make up for some if not all of the remaining 20%. To explore the contribution of another channel to survival, van den Berg et al. also queried the contribution of MscS, another mechanosensitive channel, to survival in the absence of any other species of mechansensitive channel. It was found that over the explored range of MscS channel copy numbers, the maximum survival rate was approximately 50%, suggesting that different mechanosensitive channels have an upper limit to how much protection they can confer. Both van den Berg et al. and our work show that there is still much to be learned with respect to the interplay between the various species of mechanosensitive channel as well as their regulation.

Recent work has shown that both magnitude and the rate of osmotic down shock are important factors in determining cell survival (4). In this work, we show that this finding holds true for a single species of mechanosensitive channel, even at high levels of expression. One might naïvely expect that this rate-dependent effect would disappear once a certain threshold of channels had been met. Our experiments, however, show that even at nearly 1000 channels per cell the predicted survival curves for a slow (< 1.0 Hz) and fast (≥ 1.0 Hz) are shifted relative to each other with the fast shock predicting lower rates of survival. This suggests either we have not reached this threshold in our experiments or there is more to understand about the relationship between abundance, channel species, and the shock rate.

Some experimental and theoretical treatments suggest that only a few copies of MscL or MscS should be necessary for 100% protection given our knowledge of the conductance and the maximal water flux through the channel in its open state (11, 38). However, recent proteomic studies have revealed average MscL copy numbers to be in the range of several hundred per cell, depending on the condition, as can be seen in Table 1 (15, 16, 39). Studies focusing solely on MscL have shown similar counts through quantitative Western blotting and fluorescence microscopy (3). Electrophysiology studies have told another story with copy number estimates ranging between 4 and 100 channels per cell (17, 40). These measurements, however, measure the active number of channels. The factors regulating channel activity in these experiments could be due to perturbations during the sample preparation or reflect some unknown mechanism of regulation, such as the presence or absence of interacting cofactors (41). The work described here, on the other hand, measures the *maximum* number of channels that could be active and may be able to explain why the channel abundance is higher than estimated by theoretical means. There remains much more to be leared about the regulation of activity in these systems. As the *in vivo* measurement of protein copy number becomes accessible through novel single-cell and single-molecule methods, we will continue to collect more facts about this fascinating system and hopefully connect the molecular details of mechanosensation with perhaps the most important physiological response - life or death.

**TABLE 1.** Measured cellular copy numbers of MscL. Asterisk (*) Indicates inferred MscL channel copy number from the total number of detected MscL peptides.

| Reported channels per cell | Method | Reference |
| --- | --- | --- |
| 480 ± 103 | Western blotting | (3) |
| 560* | Ribosomal profiling | (39) |
| 331* | Mass spectrometry | (15) |
| 583* | Mass spectrometry | (16) |
| 4 - 5 | Electrophysiology | (17) |
| 10 - 100 | Electrophysiology | (13) |
| 10 - 15 | Electrophysiology | (40) |

## Materials & Methods

### Bacterial strains and growth conditions

The bacterial strains are described in Table S1. The parent strain for the mutants used in this study was MJF641 (5), a strain which had all seven mechanosensitive channels deleted. The MscL-sfGFP coding region from MLG910 (3) was integrated into MJF641 by P1 transduction, creating the strain D6LG-Tn10. Selection pressure for MscL integration was created by incorporating an osmotic shock into the transduction protocol, which favored the survival of MscL-expressing stains relative to MJF641 by ~100-fold. Screening for integration candidates was based on fluorescence expression of plated colonies. Successful integration was verified by sequencing. Attempts to transduce RBS-modified MscL-sfGFP coding regions became increasingly inefficient as the targeted expression level of MscL was reduced. This was due to the decreasing fluorescence levels and survival rates of the integration candidates. Consequently, Shine-Dalgarno sequence modifications were made by inserting DNA oligos with lambda Red-mediated homologous recombination, i.e., recombineering (42). The oligos had a designed mutation (Fig. 2) flanked by ~25 base pairs that matched the targeted MscL region (Table S2). A two-step recombineering process of selection followed by counter selection using a *tetA-sacB* gene fusion cassette (43) was chosen because of its capabilities to integrate with efficiencies comparable to P1 transduction and not leave antibiotic resistance markers or scar sequences in the final strain. To prepare the strain D6LG-Tn10 for this scheme, the Tn10 transposon containing the *tetA* gene needed to be removed to avoid interference with the *tetA-sacB* cassette. Tn10 was removed from the middle of the *ycjM* gene with the primer Tn10delR (Table S2) by recombineering, creating the strain D6LG (SD0). Counter selection against the *tetA* gene was promoted by using agar media with fusaric acid (43, 44). The *tetA-sacB* cassette was PCR amplified out of the strain XTL298 using primers MscLSPSac and MscLSPSacR (Table S2). The cassette was integrated in place of the spacer region in front of the MscL start codon of D6LG (SD0) by recombineering, creating the intermediate strain D6LTetSac. Positive selection for cassette integration was provided by agar media with tetracycline. Finally, the RBS modifying oligos were integrated into place by replacing the *tetA-sacB* cassette by recombineering. Counter selection against both *tetA* and *sacB* was ensured by using agar media with fusaric acid and sucrose (43), creating the Shine-Dalgarno mutant strains used in this work.

Strain cultures were grown in 5 mL of LB-Lennox media with antibiotic (apramycin) overnight at 37°C. The next day, 50 μL of overnight culture was inoculated into 5 mL of LB-Lenox with antibiotic and the culture was grown to 0D600nm ~0.25. Subsequently, 500 μL of that culture was inoculated into 5 mL of LB-Lennox supplemented with 500mM of NaCl and the culture was regrown to 0D600nm ~0.25. A 1 mL aliquot was taken and used to load the flow cell.

### Flow cell

All experiments were conducted in a home-made flow cell as is shown in Fig. 3A. This flow cell has two inlets which allow media of different osmolarity to be exchanged over the course of the experiment. The imaging region is approximately 10 mm wide and 100 *μm* in depth. All imaging took place within 1–2 cm of the outlet to avoid imaging cells within a non-uniform gradient of osmolarity. The interior of the flow cell was functionalized with a 1:400 dilution of polyethylamine prior to addition of cells with the excess washed away with water. A dilute cell suspension in LB Lennox with 500 mM NaCl was loaded into one inlet while the other was connected to a vial of LB medium with no NaCl. This hypotonic medium was clamped during the loading of the cells.

Once the cells had adhered to the polyethylamine coated surface, the excess cells were washed away with the 500 mM NaCl growth medium followed by a small (~20 *μ*L) air bubble. This air bubble forced the cells to lay flat against the imaging surface, improving the time-lapse imaging. Over the observation period, cells not exposed to an osmotic shock were able to grow for 4–6 divisions, showing that the flow cell does not directly impede cell growth.

### Imaging conditions

All imaging was performed in a flow cell held at 30°C on a Nikon Ti-Eclipse microscope outfitted with a Perfect Focus system enclosed in a Haison environmental chamber (approximately 1°C regulation efficiency). The microscope was equipped with a 488 nm laser excitation source (CrystaLaser) and a 520/35 laser optimized filter set (Semrock). The images were collected on an Andor Xion +897 EMCCD camera and all microscope and acquisition operations were controlled via the open source μManager microscope control software (27). Once cells were securely mounted onto the surface of the glass coverslip, between 15 and 20 positions containing 5 to 10 cells were marked and the coordinates recorded. At each position, a phase contrast and GFP fluorescence image was acquired for segmentation and subsequent measurement of channel copy number. To perform the osmotic shock, LB media containing no NaCl was pulled into the flow cell through a syringe pump. To monitor the media exchange, both the high salt and no salt LB media were supplemented with a low-affinity version of the calcium-sensitive dye Rhod-2 (250 nM; TEF Labs) which fluoresces when bound to Ca^2+^. The no salt medium was also supplemented with 1*μ*M CaCl_2_ to make the media mildly fluorescent and the exchange rate was calculated by measuring the fluorescence increase across an illuminated section of one of the positions. These images were collected in real time for the duration of the shock. The difference in measured fluorescence between the pre-shock images and those at the end of the shock set the scale of a 500 mM NaCl down shock. The rate was calculated by fitting a line to the middle region of this trace. Further details regarding this procedure can be found in Bialecka-Fornal, Lee, and Phillips, 2015 (4).

### Image Processing

Images were processed using a combination of automated and manual methods. First, expression of MscL was measured via segmenting individual cells or small clusters of cells in phase contrast and computing the mean pixel value of the fluorescence image for each segmented object. The fluorescence images were passed through several filtering operations which reduced high-frequency noise as well as corrected for uneven illumination of the excitation wavelength.

Survival or death classification was performed manually using the CellProfiler plugin for ImageJ software (NIH). A survivor was defined as a cell which was able to undergo two division events after the osmotic down shock. Cells which detached from the surface during the post-shock growth phase or those which became indistinguishable from other cells due to clustering were not counted as survival or death and were removed from the dataset completely. A region of the cell was manually marked with 1.0 (survival) or 0.0 (death) by clicking on the image. The `xy` coordinates of the click as well as the assigned value were saved as an `.xml` file for that position.

The connection between the segmented cells and their corresponding manual markers was automated. As the manual markings were made on the first phase contrast image after the osmotic shock, small shifts in the positions of the cell made one-to-one mapping with the segmentation mask non-trivial. The linkages between segmented cell and manual marker were made by computing all pairwise distances between the manual marker and the segmented cell centroid, taking the shortest distance as the true pairing. The linkages were then inspected manually and incorrect mappings were corrected as necessary.

All relevant statistics about the segmented objects as well as the sample identity, date of acquisition, osmotic shock rate, and camera exposure time were saved as `.csv` files for each individual experiment. A more in-depth description of the segmentation procedure as well as the relevant code can be accessed as a Jupyter Notebook at (http://rpgroup.caltech.edu/mscl_survival).

### Calculation of effective channel copy number

To compute the MscL channel copy number, we relied on measuring the fluorescence level of a bacterial strain in which the mean MscL channel copy number was known via fluorescence microscopy (3). *E. coli* strain MLG910, which expresses the MscL-sfGFP fusion protein from the wild-type SD sequence, was grown under identical conditions to those described in Bialecka-Fornal et al. 2015 in M9 minimal medium supplemented with 0.5% glucose to an OD_600nm_ of ~0.3. The cells were then diluted ten fold and immobilized on a rigid 2% agarose substrate and placed onto a glass bottom petri dish and imaged in the same conditions as described previously.

Images were taken of six biological replicates of MLG910 and were processed identically to those in the osmotic shock experiments. A calibration factor between the average cell fluorescence level and mean MscL copy number was then computed. We assumed that all measured fluorescence (after filtering and background subtraction) was derived from the MscL-sfGFP fusion,

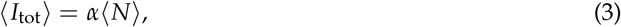

in which *α* is the calibration factor and 〈*N*〉 is the mean cellular MscL-sfGFP copy number as reported in Bialecka-Fornal et al. 2012 (3). To correct for errors in segmentation, the intensity was computed as an areal density 〈*I_A_*〉 and was multiplied by the average cell area 〈*A*〉 of the population. The calibration factor was therefore computed as

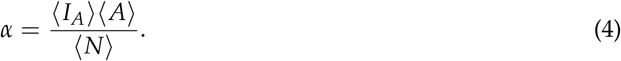

We used Bayesian inferential methods to compute this calibration factor taking measurement error and replicate-to-replicate variation into account. The resulting average cell area and calibration factor was used to convert the measured cell intensities from the osmotic shock experiments to cell copy number.

The details of this inference are described in depth in the supplemental information (*Standard Candle Calibration*).

### Logistic regression

We used Bayesian inferential methods to find the most probable values of the coefficients *β*_0_ and *β*_1_ and the appropriate credible regions and is described in detail in the supplemental information (*Logistic Regression*). Briefly, we used Markov chain Monte Carlo (MCMC) to sample from the log posterior distribution and took the most probable value as the mean of the samples for each parameter. The MCMC was performed using the Stan probabilistic programming language (45) and all models can be found on the GitHub repository (http://github.com/rpgroup-pboc/mscl_survival).

### Calculation of survival probability error

The vertical error bars for the points shown in Fig. 5 represent our uncertainty in the survival probability given our measurement of *n* survivors out of a total *N* single-cell measurements. The probability distribution of the survival probability *p_s_* given these measurements can be written using Bayes’ theorem as

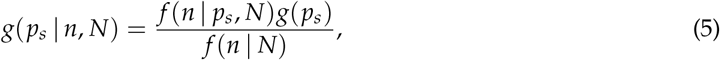

where *g* and *f* represent probability density functions over parameters and data, respectively. The likelihood *f* (*n*|*p_s_, N*) represents the probability of measuring *n* survival events, given a total of *N* measurements each with a probability of survival *p_s_*. This matches the story for the Binomial distribution and can be written as

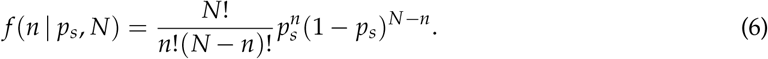

To maintain maximal ignorance we can assume that any value for *p_s_* is valid, such that is in the range [0, 1]. This prior knowledge, represented by *g*(*p_s_*), can be written as

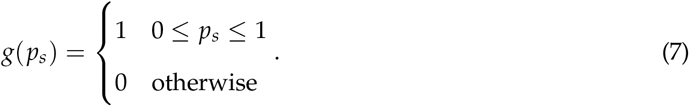

We can also assume maximal ignorance for the total number of survival events we could measure given *N* observations, *f* (*n* | *N*). Assuming all observations are equally likely, this can be written as

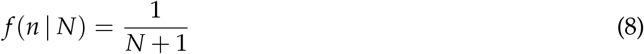

where the addition of one comes from the possibility of observing zero survival events. Combining Eqns. 6, 7, 8, the posterior distribution *g*(*p_s_| n, N*) is

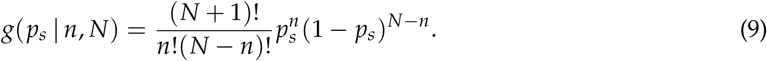

The most probable value of *p_s_*, where the posterior probability distribution given by Eq. 9 is maximized, can be found by computing the point at which derivative of the log posterior with respect to *p_s_* goes to zero,

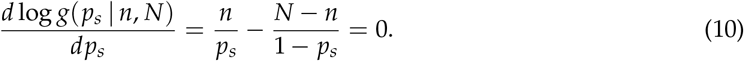

Solving Eq. 10 for *p_s_* gives the most likely value for the probability,

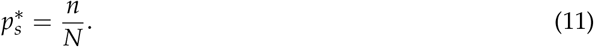

So long as 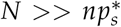, Eq. 9 can be approximated as a Gaussian distribution with a mean 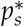 and a variance 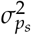. By definition, the variance of a Gaussian distribution is computed as the negative reciprocal of the second derivative of the log posterior evaluated at 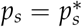,

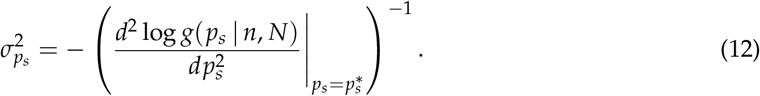

Evaluating Eq. 12 yields

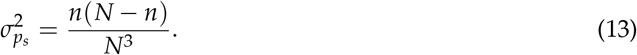

Given Eq. 11 and Eq. 13, the most-likely survival probability and estimate of the uncertainty can be expressed as

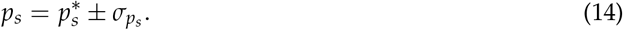

### Data and software availability

All raw image data is freely available and is stored on the CaltechDATA Research Data Repository (46). The raw Markov chain Monte Carlo samples are stored as `.csv` files on CaltechDATA (47). All processed experimental data, Python, and Stan code used in this work are freely available through our GitHub repository (http://github.com/rpgroup-pboc/mscl_survival)(48) accessible through DOI: 10.5281/zenodo.1252524. The scientific community is invited to fork our repository and open constructive issues.

## Acknowledgements

We thank Nathan Belliveau, Maja Bialecka-Fornal, Justin Bois, Soichi Hirokawa, Jaspar Landman, Manuel Razo-Mejia, Muir Morrison, and Shyam Saladi for useful advice and discussion. We thank Don Court for strain XTL298 and Samantha Miller for strain MJF641. This work was supported by the National Institutes of Health DP1 0D000217 (Director’s Pioneer Award), R01 GM085286, GM084211-A1, GM118043-01, and La Fondation Pierre Gilles de Gennes.

